# Rapid Single Cell Evaluation of Human Disease and Disorder Targets Using REVEAL: SingleCell^™^

**DOI:** 10.1101/2020.06.24.169730

**Authors:** Namit Kumar, Ryan Golhar, Kriti Sen Sharma, James L Holloway, Srikant Sarangi, Isaac Neuhaus, Alice M. Walsh, Zachary W. Pitluk

**Author notes:** equal contribution (Namit Kumar, Ryan Golhar, Kriti Sen Sharma).

## Abstract

Single-cell (sc) sequencing performs unbiased profiling of individual cells and enables evaluation of less prevalent cellular populations, often missed using bulk sequencing. However, the scale and the complexity of the sc datasets poses a great challenge in its utility and this problem is further exacerbated when working with larger datasets typically generated by consortium efforts. As the scale of single cell datasets continues to increase exponentially, there is an unmet technological need to develop database platforms that can evaluate key biological hypothesis by querying extensive single-cell datasets.

Large single-cell datasets like human cell atlas and COVID-19 cell atlas (collection of annotated sc datasets from various human organs) are excellent resources for profiling target genes involved in human diseases and disorders ranging from oncology, auto-immunity, as well as infectious diseases like COVID-19 caused by SARS-CoV-2 virus. SARS-CoV-2 infections have led to a worldwide pandemic with massive loss of lives, infections exceeding 7 million cases. The virus uses ACE2 and TMPRSS2 as key viral entry associated proteins expressed in human cells for infections. Evaluating the expression profile of key genes in large single-cell datasets can facilitate testing for diagnostics, therapeutics and vaccine targets; as the world struggles to cope with the on-going spread of COVID-19 infections.

In this manuscript we describe, REVEAL: SingleCell which enables storage, retrieval and rapid query of single-cell datasets inclusive of millions of cells. The analytical database described here enables selecting and analyzing cells across multiple studies. Cells can be selected using individual metadata tags, more complex hierarchical ontology filtering, and gene expression threshold ranges, including co-expression of multiple genes. The tags on selected cells can be further evaluated for testing biological hypothesis. One such example includes identifying the most prevalent cell type annotation tag on returned cells.

We used REVEAL: SingleCell to evaluate expression of key SARS-CoV-2 entry associated genes, and queried the current database (2.2 Million cells, 32 projects) to obtain the results in <60 seconds. We highlighted cells expressing COVID-19 associated genes are expressed on multiple tissue types, thus in part explains the multi-organ involvement in infected patients observed worldwide during the on-going COVID-19 pandemic.

## Background

Single cell RNA sequencing (scRNAseq) datasets have played a crucial role in identifying specific cell types in airway tissues that express the SARS-CoV-2 virus receptor, ACE2, and host responses in peripheral blood(1). With more than 7 million cases of SARS-CoV-2 infection (COVID-19) and 403,000 fatalities reported world-wide (8 June 2020)(2), SARS-CoV-2 interventions are an unmet medical need of pandemic proportions(3, 4). Rapid identification of cell-type-specific expression and co-expression of the targets can identify novel cellular subtypes(5), facilitate decisions about biomarkers for target engagement(6) and response(7), potential delivery methods for therapies, and detection methods for diagnosis(8). Additional host factors, TMPRSS2 and Cathepsin B/L, play a key role in the virus infection process and may be used as biomarkers and/or drug targets alone or in combination with ACE2. Peripheral responses may include the appearance of novel immune cellular subtypes and the absence of overexpression of traditional cytokine storm peptides(9). COVID interactome map(10) serves as a rich resource set of approved medicines for testing once the tissue abundance is confirmed in COVID-19 patients.

While the field of precision medicine has steadily advanced through the elucidation of bulk tissue or fluid biomarkers, there is exciting potential for new discoveries due to scRNAseq. scRNAseq is capable of identifying rare cell populations or markers on cellular subsets, associating cellular subsets with disease onset and/or treatment response. Single cell data collections like the COVID-19 Cell Atlas(11) (CCA) and the Human Cell Atlas(12) (HCA) are resources for expression profiling of key targets involved in SARS-CoV-2 infection of the cells and the subsequent immune response. However, the full utility of these data collections is limited due to a lack of database management strategy that allows facile cross comparison of the distribution and levels of specific gene expression between samples and projects without a significant bioinformatics and computational effort. For instance, determining the tissue distribution of expressed targets can enable rapid decisions for drug delivery methods and potential combination therapies. Without new data solutions, simple queries can become lengthy processes due to the scale of the datasets as well as the programming and computational resources required.

Ease of accessing and evaluating multiple scRNAseq data sets for the purposes of developing better therapeutic targets and biomarkers for clinical studies presents a fundamental challenge for their use in precision medicine. Seyhan et al. suggested that an important milestone for implementing precision medicine will be creating an “accessible data commons” to streamline biomarker discovery and simplify tests for the mechanism of action.(13) For the authors, the term accessible means easily searched by non-programmer biomedical scientists for subsets of relevant data. The challenge is creating a data management and analysis capability that facilitates the comparison of small diseased tissue datasets, collected in the clinic, to other diseased tissue datasets in the pubic domain as well as to large healthy tissue datasets, like the Human Cell Atlas (HCA)(12). These comparisons may identify the presence or emergence of subpopulations of cells that are resistant to therapy, or they could indicate infiltration or other cellular changes that would be elusive in either bulk RNAseq experiments or in flow cytometry, which are limited in the number of markers monitored.(14) The need for potentially high numbers of biological replicates to identify differential gene expression (DGE) will only accentuate the need for a data commons.(15, 16)

This study describes the scalable REVEAL: SingleCell platform developed to address the issue of enabling rapid queries of multiple large single cell datasets, like the HCA, on the order of millions of cells. This study represents the first phase of a project to develop the framework necessary for searching across, analyzing, and in the future, implementing machine-learning in a data commons comprised of single cell precision medicine data sets. REVEAL: SingleCell addresses the challenge of storing large sparse arrays from various studies in a FAIR (findable, accesible, interoperable, reusable) manner. REVEAL: SingleCell is built on top of SciDB, an array native computational database that has R, Python, and REST APIs(17).

We loaded normalized scRNAseq data into the REVEAL: SingleCell platform to allow searching across reference datasets to find the distribution of transcripts for ACE2, TMPRSS2 and other host factors. The same schema and commands can be adapted for use with other single cell ‘omics data such as CITE-seq, snRNAseq and other data types. We provide timings for retrieving data that highlight the time challenges of the repetitive ETL (extract, transform and load) process that workflows like the Seurat(18) and HCAData(19) packages present.

## Construction and Content

### Construction

Single cell data sets are loaded into SciDB, a unified scientific data management and computational platform organized around vectors and multi-dimensional arrays as the basic data modeling, storage, and computational unit.(20) The data model accommodates rapid and FAIR access to heterogeneous, multi-attribute data as well as metadata like ontologies and reference data sets. Multiple users can load, read, and write data in a secure, transactionally safe manner as data operations are guaranteed to be atomic and consistent (ACID compliant). The REVEAL: SingleCell solution is an app built on top of SciDB that provides purpose-built data schema, interfaces, and task-focused functionality, using controlled vocabulary. A Shiny GUI supports data visualization and exploration by non-programming scientists. R and Python APIs provide direct, *ad hoc* access and analysis, as well as extensibility via the integration of additional library packages. A FLASK(17) REST API implements a web interface. Documentation is provided as R markdown notebooks along with context-sensitive online help. Figure 1 provides a detailed view of the APIs, security, and storage architecture for SciDB implemented on AWS.

**Figure 1:**
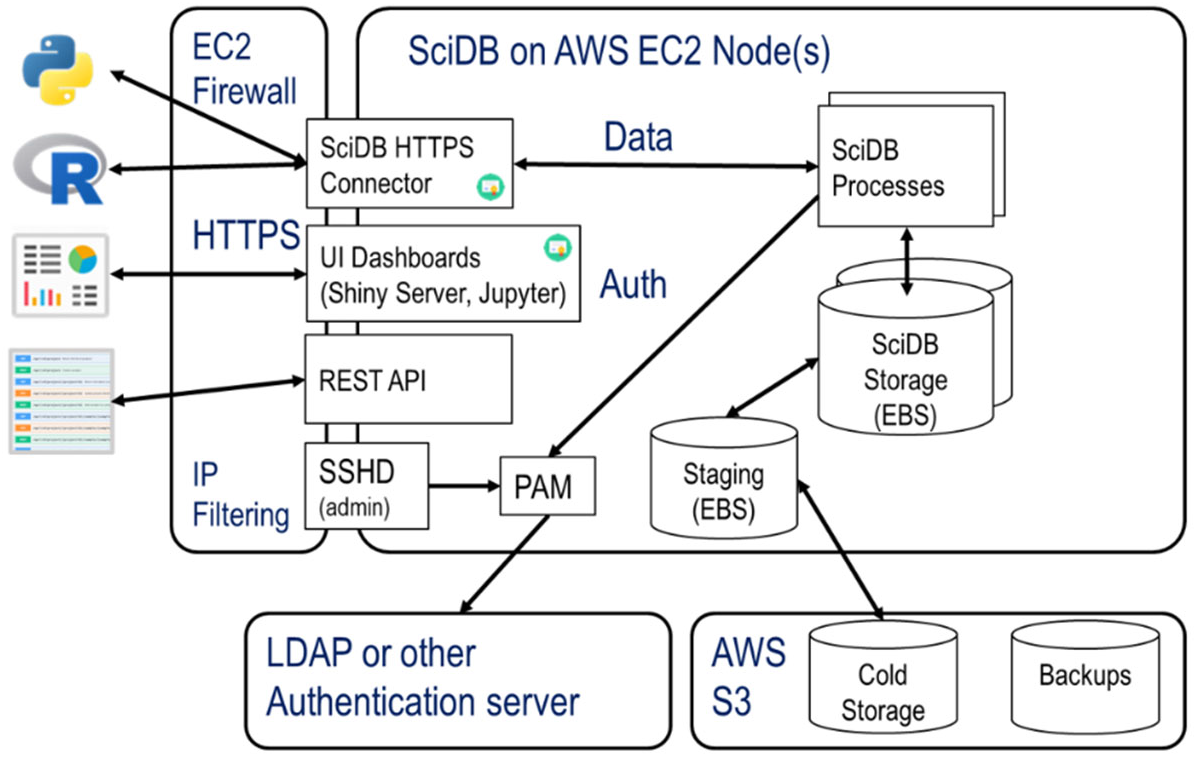
system configuration. Figure Legend REVEAL: SingleCell implementation in EC2: SciDB offers multiple paths to retrieve and load data. There are REST, R and Python APIs for server-side communication, R can also communicate via a local machine using HTTPS. The data and transactions are all ACID compliant. In this EC2 instance of REVEAL for scRNAseq, a 16-core machine with 64 GB of RAM, and 500 GB of SSD is used.

The software versions used are shown below in Table 1.

**Table 1:** Software requirements. Table 1 Legend: a list of software versions used for analysis

|  |  |  |
| --- | --- | --- |
| **Linux** | CentOS 7.5 /<br>Ubuntu 14.04 |  |
| **SciDB** | 19.11.5 |  |
| **R** | 3.6 | https://www.r-project.org |
| **R packages** |  |  |
| - Seurat | 3.1.x | https://satijalab.org/seurat/ |
| - SciDBR | 2.0.2 | github.com/Paradigm4/scidbr |
| - revealgenomics | 0.6 | private github |
| - revealsc | 0.1.0 | private github |
| **Python** | 3.7.6 | https://www.python.org |
| **Python REST API** | 1.1.2 | https://flask.palletsprojects.com/en/1.1.x/ |

### Content

The following publicly available datasets were loaded: Human Cell Atlas (HCA) Census of Immune Cells data set(21), COVID-19 Cell Atlas (CCA)(11) (excluding the Aldinger, *et al*. Fetal Cerebellum data set). These datasets were all aligned to the GRCh38 reference genome. Data sizes into the hundreds of TBs are feasible. The current system contains 32 projects, totalling less than 1 TB.

HCA provided filtered raw counts data in 10x CellRanger version 3.0 format. This data was loaded into R as a Seurat object, normalized using the Seurat scTransform algorithm(22) and then converted back to 10x CellRanger v3.0 format. The CCA provided normalized data in .h5ad format as used in the Python Scanpy(23) and anndata(24) libraries. CCA .h5ad files were converted into the 10x CellRanger format (using standard convertors from the Python anndata, scipy.io(25) libraries). In both cases, the cell metadata tags (e.g., CellType, percent.mt) were saved as .tsv files from the normalized Seurat object (HCA) and .h5ad files (CCA), and loaded into the database using the REST API metadata update endpoints. The REST API checked for consistency in the 10x format, missing values, among others.

### Content schema

Data are modelled as multi-dimensional arrays. Each element in an array contains one or more attributes. Modelling data as arrays enables rapid sub-setting of cells by gene expression levels, ontology and QC tags, individually and in combination across samples.

Figure 2 illustrates the various single cell data submodalities that can also be stored in the array elements of the n-dimensional SciDB arrays. Though this project stored only scRNAseq data, the multi-dimensional array schema can be extended to hold many complimentary data types, including snRNAseq, scATAC-seq, CITE-seq, among others.

**Figure 2:**
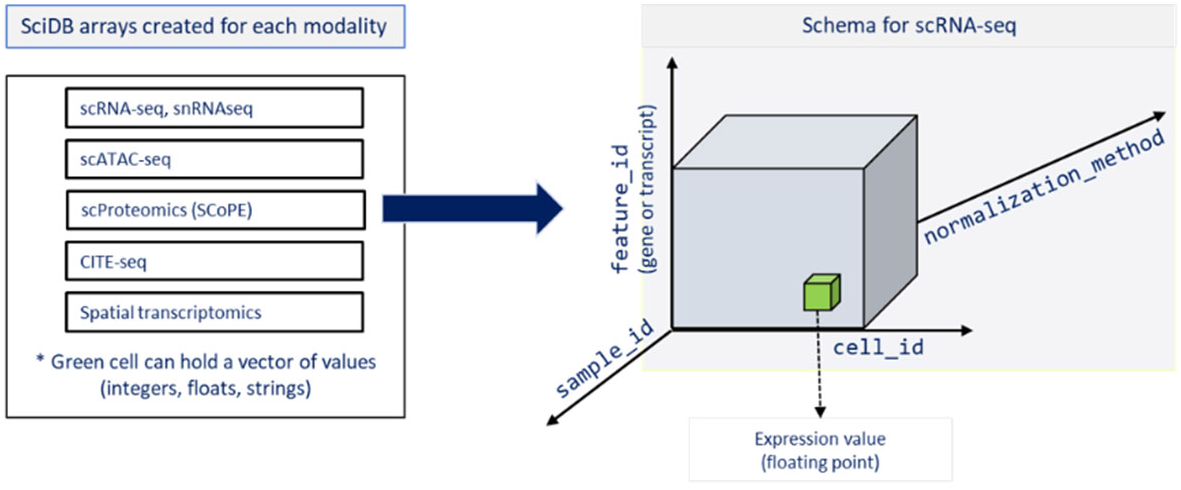
single cell data types compatible with REVEAL: SingleCell. Figure 2 Legend: single cell data types compatible with REVEAL: SingleCell. Elements in the n-dimensional arrays can contain several orthogonal omics data types, as mentioned in the figure.

### Content data hierarchy

Figure 3 illustrates the hierarchical relationship of metadata. The label “projects” was used for collections of samples which are often also referred to as studies. For instance, the HCA Census of Immune Cells is one project with 16 samples. At the sample level, anatomy/tissue type and disease type are selectable with the UBERON and DOID identifiers. At the cell level, CL IDs were used to enable selection of specific cell types. It is important to note that there was tremendous heterogeneity in how the information was presented in these individual projects, and an automated system for unification is being developed. Feature sets(26) include information about the human genome version and the sub-category feature, allowing selection by either ENSEMBL ID or gene symbol. Gene symbols were used because most public data are not annotated with ENSEMBL IDs. Due to the diversity of the metadata (especially when sourced from public studies), we stored metadata as key-value pairs in the elements of the sample array shown below in Table 2.

**Figure 3:**
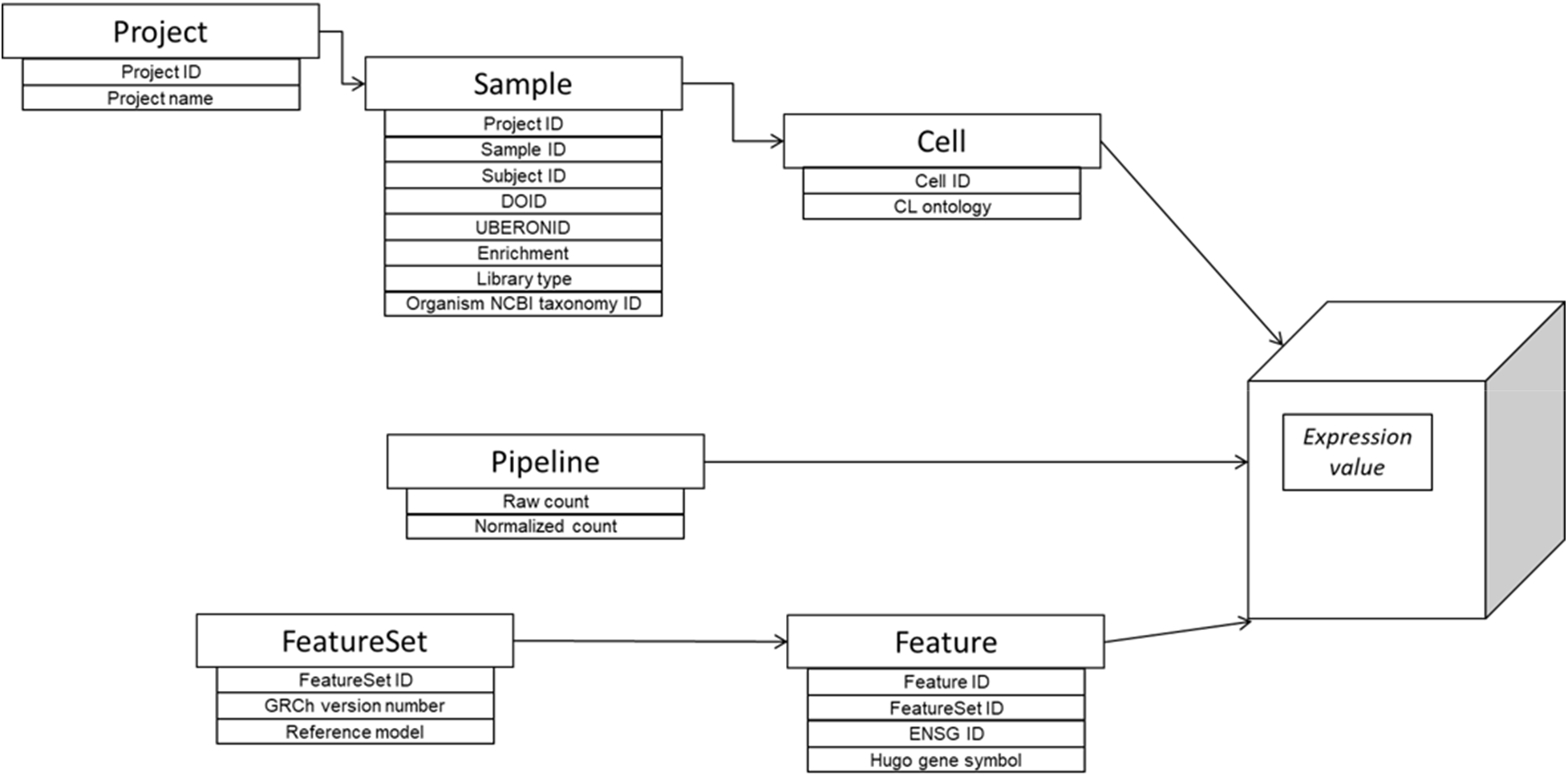
data hierarchy.

**Table 2.**
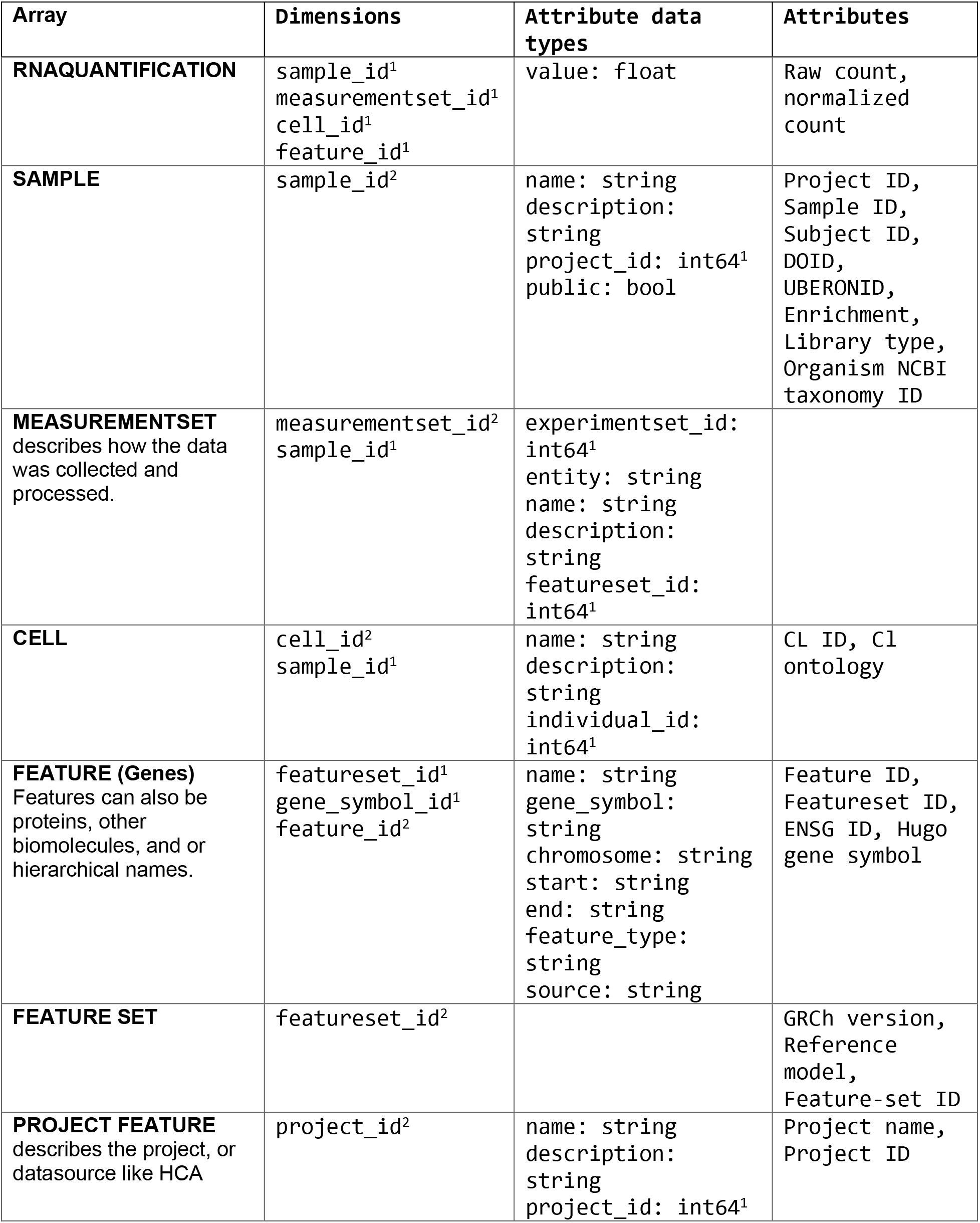
Arrays and attributes in REVEAL: SingleCell. Table 2 Legend: Table 2 shows the schema. Data of interest can be accessed and filtered by their dimensions and attributes. The superscript 1 indicates primary dimensions for selection, and the superscript 2 inidcates secondary dimensions for selection. The general categories for attributes include but are not limited to:

| Array | Dimensions | Attribute data types | Attributes |
| --- | --- | --- | --- |
| **RNAQUANTIFICATION** | sample_id<sup>1</sup><br>measurementset_id<sup>1</sup><br>cell_id<sup>1</sup><br>feature_id<sup>1</sup> | value: float | Raw count,<br>normalized<br>count |
| **SAMPLE** | sample_id<sup>2</sup> | name: string<br>description:<br>string<br>project_id: int64<sup>1</sup><br>public: bool | Project ID,<br>Sample ID,<br>Subject ID,<br>DOID,<br>UBERONID,<br>Enrichment,<br>Library type,<br>Organism NCBI<br>taxonomy ID |
| **MEASUREMENTSET**<br>describes how the data<br>was collected and<br>processed. | measurementset_id<sup>2</sup><br>sample_id<sup>1</sup> | experimentset_id:<br>int64<sup>1</sup><br>entity: string<br>name: string<br>description:<br>string<br>featureset_id:<br>int64<sup>1</sup> |  |
| **CELL** | cell_id<sup>2</sup><br>sample_id<sup>1</sup> | name: string<br>description:<br>string<br>individual_id:<br>int64<sup>1</sup> | CL ID, Cl<br>ontology |
| **FEATURE (Genes)**<br>Features can also be<br>proteins, other<br>biomolecules, and or<br>hierarchical names. | featureset_id<sup>1</sup><br>gene_symbol_id<sup>1</sup><br>feature_id<sup>2</sup> | name: string<br>gene_symbol:<br>string<br>chromosome: string<br>start: string<br>end: string<br>feature_type:<br>string<br>source: string | Feature ID,<br>Featureset ID,<br>ENSG ID, Hugo<br>gene symbol |
| **FEATURE SET** | featureset_id<sup>2</sup> |  | GRCh version,<br>Reference<br>model,<br>Feature-set ID |
| **PROJECT FEATURE**<br>describes the project, or<br>datasource like HCA | project_id<sup>2</sup> | name: string<br>description:<br>string<br>project_id: int64<sup>1</sup> | Project name,<br>Project ID |

■ scRNAseq expression values, both normalized and raw counts
■ categorical and continuous tags which can contain metadata on any entities from the pipeline used to generate the tags.

- projects, e.g. data generation source (public, institutional-internal)
- samples, e.g. UBERONID; DOID; organ (lung, rectum, illium)
- cells, e.g. CL ID; cell type (CD8+, enterocytes); percent.mt (percent mitochondria)
- features, e.g. strand (+, -); biotype (protein-coding, frameshift) Note that the tags, UBERONID, DOID, and CL ID, hold controlled vocabulary from publicly curated ontologies like Ontobee. These tags enable hierarchical searches, e.g. search for all cells matching CLID CL:0000584 (enterocyte) and its children.

### Content data curation

Cell type is one of the most important selection criteria. However public datasets in CCA used multiple disparate naming conventions, e.g. cell_type, CellType, celltypes, celltype1. These names were retained as is in the database, but an extra tag, CellType.select, was added for harmonization across all projects. The CellType.select tag was manually curated.

Subject-level and sample-level metadata were often missing in the CCA. We provide a manually curated supplementary table with the exact numbers of subjects and samples, where it was possible to obtain the information (S1).

### Queries and REST API

Table 3 lists the queries and functions built into the REVEAL: SingleCell app.

**Table 3:**
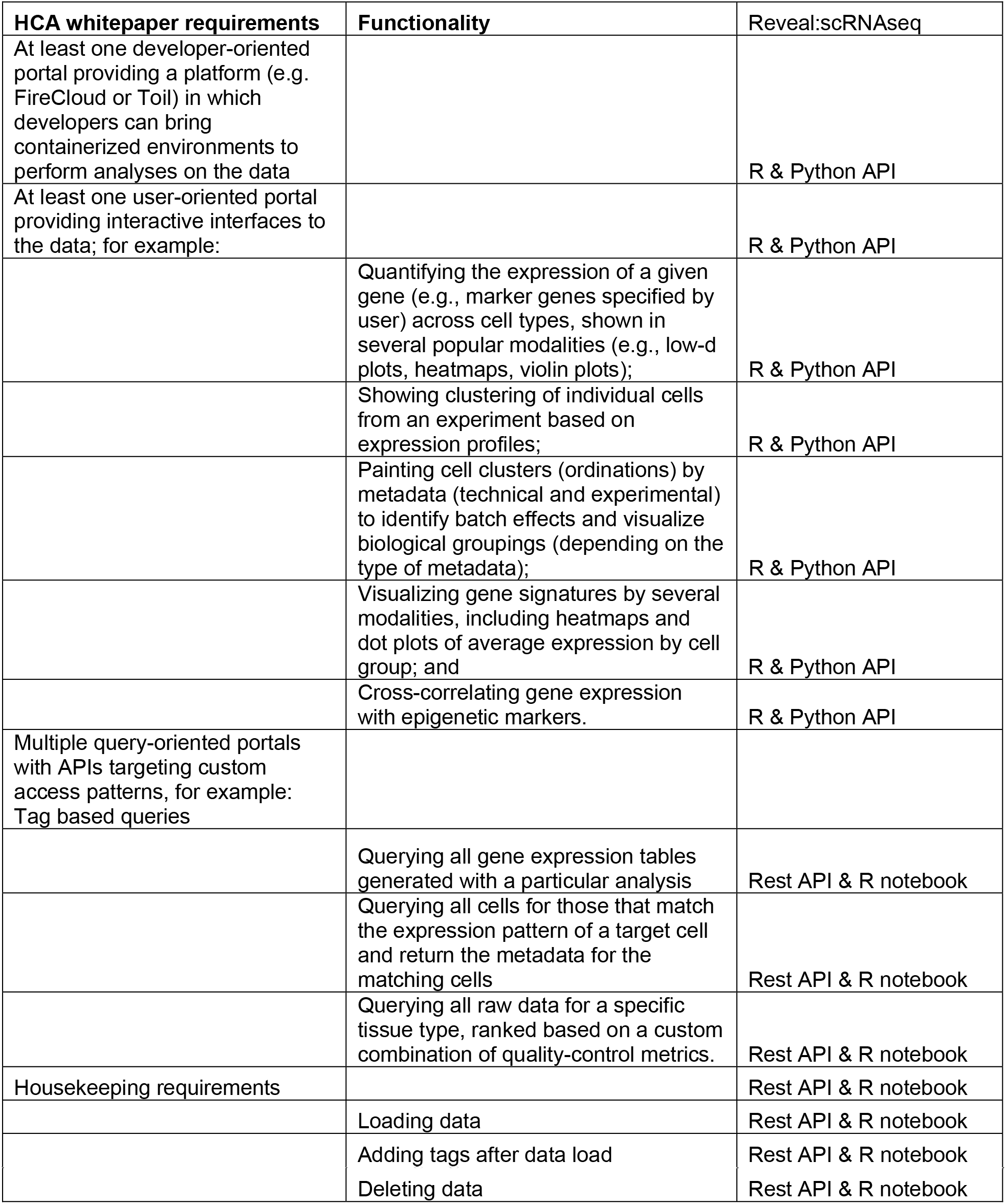
Queries and functions built into the database. Table 3 legend: The requirements listed in the HCA whitepaper take two forms: actual queries and visualization capabilities. The R and Python APIs support the visualization requirements. The REST API and R notebook support the queries. We included the housekeeping requirements in the list because those are essential capabilities for a database. These are accessible through an R API and REST API.

**Figure 4.**
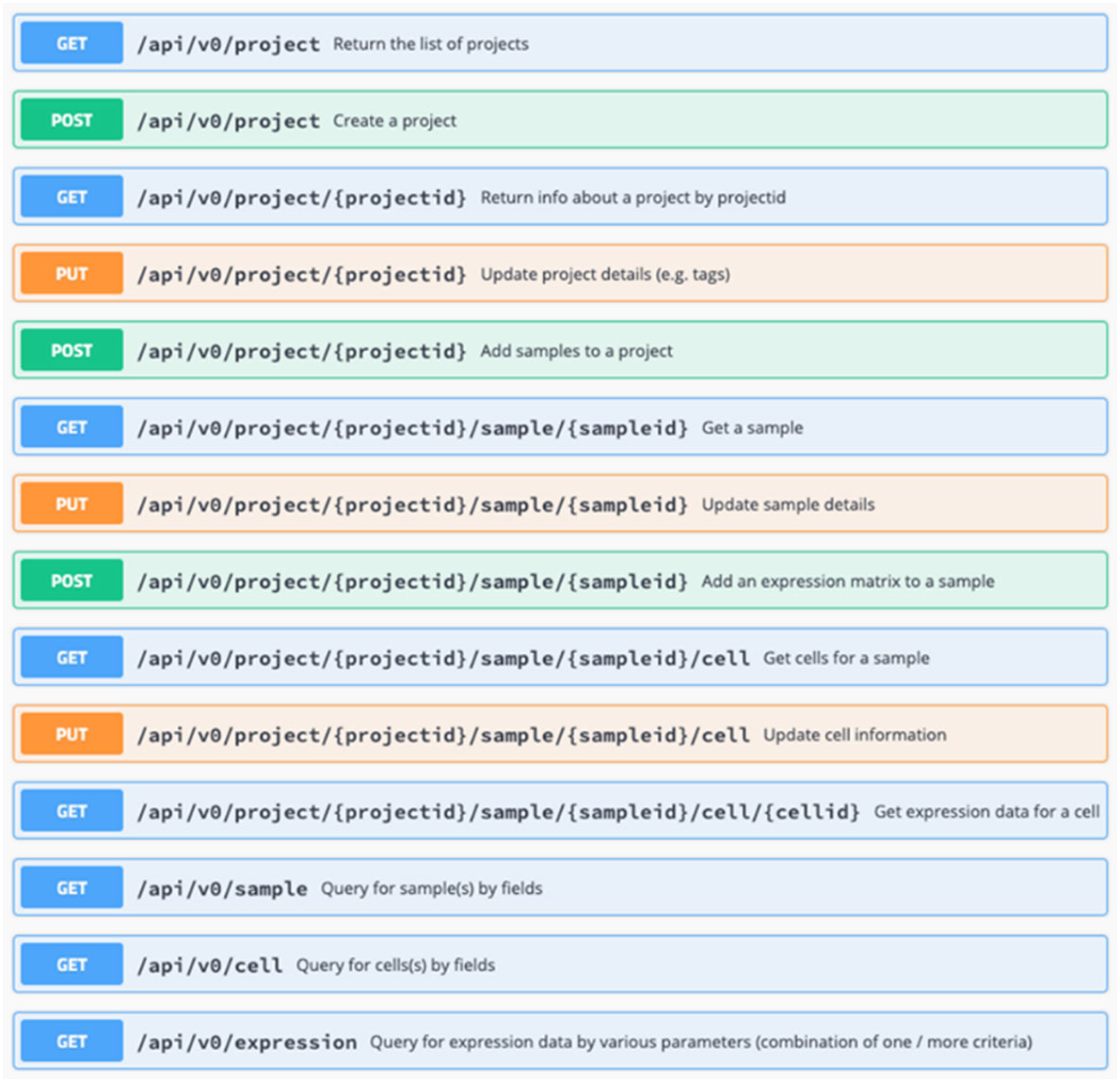
lists the REST API commands.

## Utility and Discussion

We approached the challenges of creating a data commons by deploying a scientific computational database, SciDB. There are distinct benefits to having scRNAseq data organized as arrays in a database, such as allowing cross-study selection of cells by gene expression thresholds or metadata tags and analysis by multiple users, while ensuring the consistency from a shared version of QA’d data and workflows. SciDB endows REVEAL: SingleCell with future-readiness, the capability of integrating genomic, proteomic, image and metabolomic data types into the same database, enabling a data commons.

Many researchers use Seurat objects or HDF5 files for storage of both scRNAseq data and calculated results. This approach contradicts the basic concept of FAIR data because each object is a silo of data. Cross-study analysis with Seurat requires loading the studies of interest into a single Seurat object and repeating a Seurat object merge step for each desired set of studies and is often limited by RAM. Thus, analysis is limited by the size of the compute hardware, i.e. RAM, to fewer than 1 million cells. Yet, the outlook is for dataset sizes to grow especially when coupled with flow cytometry, microscopy and new methods. For example, single cell and single nucleus data sets range in complexity from analysis of total mRNAs, to capped RNAs to transcriptional velocity to transient physiologic responses(27), many of which may be inter-compared to test hypotheses.(28) Emerging higher throughput and lower cost methods of single cell transcriptional profiles like Sci-Plex, will create much larger data sets to search across and analyze.(29)

REVEAL: SingleCell was designed as a data commons with the goal of removing silos, supporting cross-study analysis, and enabling scaling of computation beyond a single instance. We populated the REVEAL: SingleCell platform with scRNAseq data from the HCA and CCA (content and construction). The same schema and commands can be used with other single cell ‘omics data such as CITE-seq(30) and snRNAseq data(31).

As a design guide, we implemented the requirements for querying data outlined in the HCA whitepaper. The HCA whitepaper didn’t include provisions for an actual database; storage was based on file retrieval. The requirements for precision medicine put a premium on being able to inter-compare datasets without needing increasingly larger amounts of RAM.

■ Querying all gene expression data generated with a particular analysis,
■ Querying all cells for those that match the expression pattern of a target cell and return the metadata for the matching cells; and
■ Querying all raw data for a specific tissue type, ranked based on a custom combination of quality-control metrics.

Table 2 shows the schema, a collection of 7 arrays. This schema fulfills the requirements for queries laid out in Table 3, allowing sub-setting of cells by gene expression levels, ontology and QC tags, individually and in combination across samples. Using the REVEAL: SingleCell platform, more complex queries relating to ontologies as well as to gene expression levels (or other continuous variables like x, y coordinates or time), or patterns can be combined. This is enabled because each element in an n-dimensional SciDB array can have unlimited numbers of tags that can be used for selection (Table 2, Figure 3). Thus, users can:

■ Query for gene expression in cells matching a cell type, and then expand the search to include cell types that are parents or children in a cell type ontology.
■ Query for gene expression to return cells with gene expression above, below, or within thresholds (e.g., ACE2 >3, <7, 4-6).
■ Query raw and/or normalized counts for each cell.

## Applying REVEAL: SingleCell to evaluate key regulators involved in SARS-CoV-2 infection

In this early phase of SARS-CoV-2 research, hypotheses regarding tissue/cell type distribution of host cofactors for viral infection (receptors, processing enzymes) and pathogenesis (changes in normal cell gene expression profiles) need to be tested quickly. As an illustration of the capabilities of REVEAL: SingleCell, we queried for all cells in the database (datasets from CCA, HCA) that either express the receptor for SARS-CoV-2, *ACE2*, the cell surface receptor for SARS-CoV-2(32), and entry facilitating enzyme, transmemembrane serine protease, *TMPRSS2*(33), or coexpress both mRNAs with *DPP4*, the receptor for MERS-CoV(34) (Table 4, 5, and Figure 5). An example of a more complex query is shown (Table 4, query 6): sequentially applying a metadata filter and then a gene expression filter on the results. These findings highlight that REVEAL: SingleCell returned results that can support interactive hypothesis generation and testing by searching across more than 30 datasets in a timespan of seconds.

**Figure 5.**
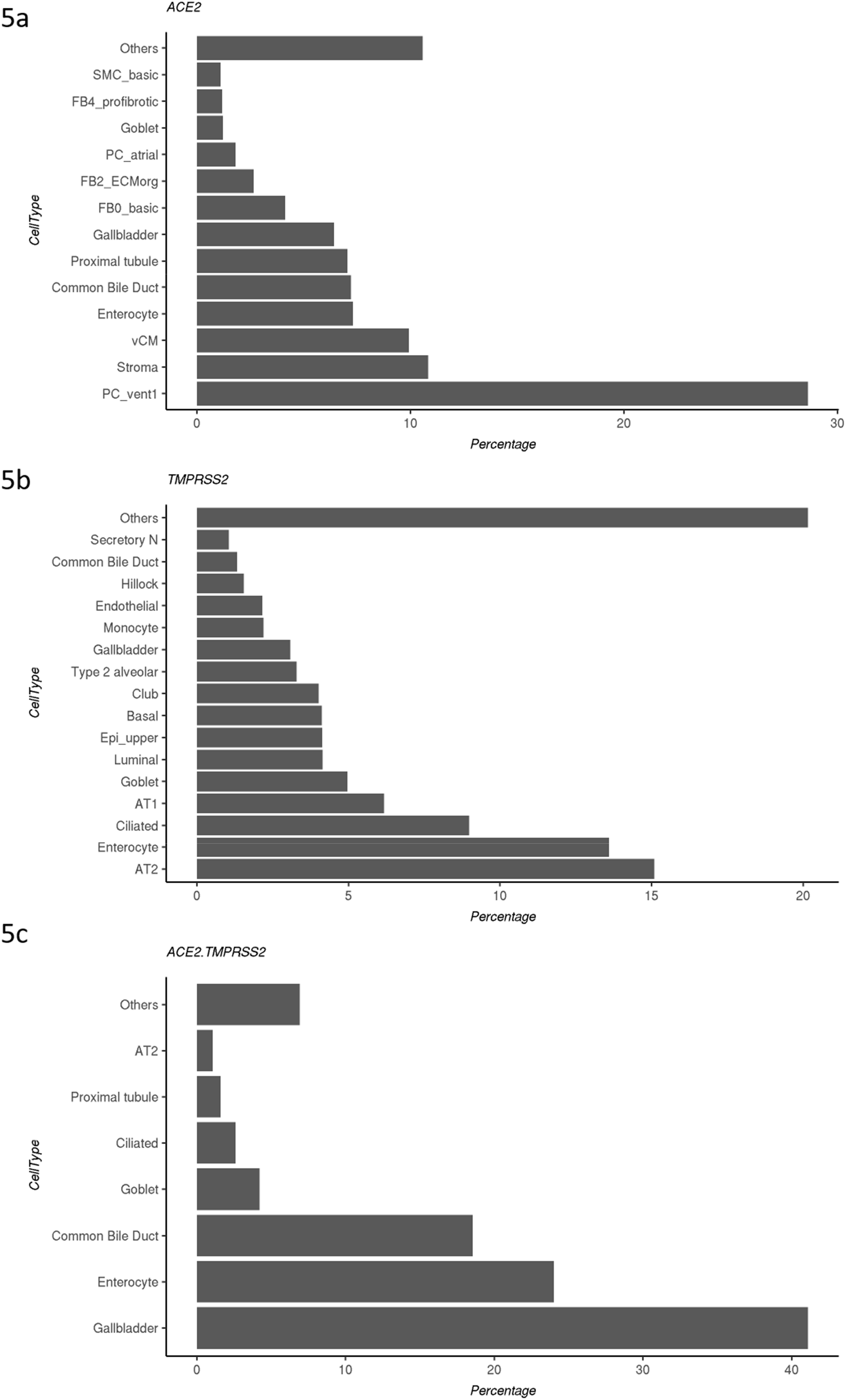
Cells matching search criteria are grouped by their cell type annotation. Cell types tags with < 1 % of total cells matching search criteria, were grouped together as ‘Others’. For co-expression, the same cell is required to express both the genes above the set thresholds.

**TABLE 4:** Benchmarking queries. Table 4 legend Queries were organized as: searching by metadata tags (1 & 2), searching by co-expression (3, 4), searching by gene list (5), searching the results of query 1 by expression levels (6), and returning the results of a project.

| Capabilities | Search criteria | Query # | Search result<br># of cells returned<br>(# of projects with data) | Total time<br>(sec) |
| --- | --- | --- | --- | --- |
| **Selecting a subset of cells (searching across 2.2M cells and 33 projects)** |  |  |  |  |
| By tags | 1 tag<br>CellType.select is any of:<br>['Enterocyte', 'Enterocytes',<br>'Best4+ Enterocytes', 'Enterocyte<br>Progenitors', 'Immature Enterocytes<br>1', 'Immature Enterocytes 2'] | 1 | 19K cells<br>(5 projects) | 14 |
|  | 2 tags<br>Above criteria on CellType.select<br>& Location is any of:<br>['Rectum', 'Decidua', 'Ileum'] | 2 | 4827 cells<br>(2 projects) | 9 |
| **Selecting cells across projects** |  |  |  |  |
| Checking for co<br>expression in more<br>than 1 gene, when<br>expression value lies in<br>a range<br>•Threshold:<br>value >= 1<br>•Restricting<br>search to<br>normalized<br>data | 'ACE2', 'TMPRSS2' | 3 | 2282 cells<br>(21 projects) | 26 |
|  | 'ACE2', 'TMPRSS2', 'DPP4' | 4 | 561 cells<br>(11 projects) | 32 |
| **Search expression** |  |  |  |  |
| By gene list across all projects | 'ACE2', 'TMPRSS2', 'DPP4' | 5 | 225K rows<br>(32 projects;<br>download size:<br>8 MB) | 15 |
| By cells across multiple projects | Using the result of Query 1 to search expression on those cells i.e. searching by ~19,000 cells (in the projects with data) | 6 | 26.7M rows<br>(5 projects;<br>download size: 1019 MB) | 27 |
| By selected project | Project: 'wang20_rectum';<br>matrix_count: 'normalized' | 7 | 11.6M rows<br>(1 project;<br>download size:<br>621 MB) | 17 |

Table 4 lists the times to return an R data frame in RStudio from querying REVEAL: SingleCell for the listed queries across many or all of the samples from CCA and HCA.

We evaluated multiple samples from CCA and HCA to identify cell type tag of cells expressing *ACE2, TMPRSS2*, and co-expression of both the markers. All cells matching the above criteria were grouped together by their cell type tags and reported as percentage of total cells matching criteria. Cell type tags with <1 % of total cells matching criteria were grouped together and labelled as ‘Others’ and the sum of their percentanges was also reported.

Our analysis, based on all cells currently loaded in the database (Figure 5), highlights that the majority of cells expressing *ACE2* have a cell type tag of PC_vent1 (Heart tissue); the majority of cells expressing *TMPRSS2* have a cell type tag of AT2 (alveolar epithelial type II cells found in the lung parenchyma); and most cells co-expressing both *ACE2* and *TMPRSS2* are tagged as Gall bladder cells. These results are consistent with previous studies.(1, 35–37).

These results in part, explain the the multi-organ involvement in infected patients observed worldwide during the on-going COVID-19 pandemic, as multiple cell types in the human body express genes utilized by SARS-CoV-2 for infection. REVEAL: SingleCell enables quick profiling of key genes involved in the current pandemic and supports additional use cases that require evaluation across a large database of single cell expression datasets such as vaccine candidates for infectious diseases, biomarkers for oncology patient stratification, and immunology-related disorders.

## Conclusion

In this paper, we introduce the REVEAL: SingleCell database that addresses immediate needs for SARS-CoV-2 research and has the potential to be used more broadly for many precision medicine applications. We used the REVEAL: SingleCell database as a reference to ask questions relevant to drug development and precision medicine regarding cell type and coexpression for genes that encode proteins necessary for SARS-CoV-2 to enter and reproduce in cells.

### Significance

The COVID-19 atlas used in this project is an example of an extensive reference dataset that can be used for understanding individual patient responses to novel therapies relative to untreated and un-infected patient data. Implementation of REVEAL: SingleCell harnesses the power of working with large and complex single cell datasets and unlocks their potential by significantly speeding up the process of selecting and analyzing data for understanding and treating individual patients using precision medicine. The next phase of development will be to extend the REVEAL: SingleCell architecture to include additional relevant datasets, as well as include other omics data types from single cell experiments.

## Declarations

Ethics approval and consent to participate – Not applicable

Consent for publication – Not applicable

### Availability of data and materials

Data is available at these websites:
COVID-19 Cell Atlas. https://www.covid19cellatlas.org. Accessed 06 June 2020.
Human Cell Atlas Data Portal: Mapping the human body at the cellular level.
https://data.humancellatlas.org. Accessed 06 June 2020.

### Competing interests

Namit Kumar, Ryan Golhar, James L. Holloway, Brian Kidd, Alice M. Walsh, Isaac Neuhaus are employees of Bristol Myers Squibb.

Kriti S. Sharma, Srikant Sarangi and Zachary Pitluk are employees of Paradigm4, Inc.

### Funding

this project was privately funded by Bristol Myers Squibb and Paradigm4.

### Author’s contributions

N.K., R.G., J.L.H., and K.S. developed the software and testing procedures. N.K., K.S., and S.S. performed analysis. Z.W.P. researched and compiled literature resources. N.K., A.M.W., Z.W.P. drafted the manuscript with input from all authors. I.N., and Z.W.P. supervised the study. N.K., R.G., A.M.W., J.L.H., I.N., Z.W.P conceptualized and designed the study.

## Acknowledgements

We would like to acknowledge Melanie McCorry for her help in developing the ontology hierarchy search function. We acknowledge Marilyn Matz for her editing support.

## Notes

https://www.covid19cellatlas.org

https://data.humancellatlas.org

## References

1. Sungnak W, Huang N, Becavin C, Berg M, Queen R, Litvinukova M, et al. SARS-CoV-2 entry factors are highly expressed in nasal epithelial cells together with innate immune genes. Nat Med. 2020;26(5):681–7.

2. Staff R. Thompson Reuters; 2020 [Available from: www.reuters.com.

3. Zhu N, Zhang D, Wang W, Li X, Yang B, Song J, et al. A Novel Coronavirus from Patients with Pneumonia in China, 2019. N Engl J Med. 2020;382(8):727–33.

4. Organization WH. Naming the coronavirus disease (COVID-19) and the virus that causes it. [Available from: https://www.who.int/emergencies/diseases/novel-coronavirus-2019/technical-guidance/naming-the-coronavirus-disease-(covid-2019)-and-the-virus-that-causes-it.

5. Farber DL, Sims PA. Dissecting lung development and fibrosis at single-cell resolution. Genome Med. 2019;11(1):33.

6. Anand P, Puranik A, Aravamudan M, Venkatakrishnan AJ, Soundararajan V. SARS-CoV-2 strategically mimics proteolytic activation of human ENaC. Elife. 2020;9.

7. Wen W, Su W, Tang H, Le W, Zhang X, Zheng Y, et al. Immune cell profiling of COVID-19 patients in the recovery stage by single-cell sequencing. Cell Discov. 2020;6:31.

8. Bertram S, Heurich A, Lavender H, Gierer S, Danisch S, Perin P, et al. Influenza and SARS-coronavirus activating proteases TMPRSS2 and HAT are expressed at multiple sites in human respiratory and gastrointestinal tracts. PLoS One. 2012;7(4):e35876.

9. Wilk AJ, Rustagi A, Zhao NQ, Roque J, Martinez-Colon GJ, McKechnie JL, et al. A single-cell atlas of the peripheral immune response to severe COVID-19. medRxiv. 2020.

10. Gordon DE, Jang GM, Bouhaddou M, Xu J, Obernier K, White KM, et al. A SARS-CoV-2 protein interaction map reveals targets for drug repurposing. Nature. 2020.

11. Atlas C-C. COVID-19 Cell Atlas [Available from: https://www.covid19cellatlas.org.

12. Atlas HC. Human Cell Atlas Data Portal: Mapping the human body at the cellular level [Available from: https://data.humancellatlas.org.

13. Seyhan AA, Carini C. Are innovation and new technologies in precision medicine paving a new era in patients centric care? J Transl Med. 2019;17(1):114.

14. Zeggini E, Gloyn AL, Barton AC, Wain LV. Translational genomics and precision medicine: Moving from the lab to the clinic. Science. 2019;365(6460):1409–13.

15. Schurch NJ, Schofield P, Gierlinski M, Cole C, Sherstnev A, Singh V, et al. How many biological replicates are needed in an RNA-seq experiment and which differential expression tool should you use? RNA. 2016;22(6):839–51.

16. Li X, Cooper NGF, O’Toole TE, Rouchka EC. Choice of library size normalization and statistical methods for differential gene expression analysis in balanced two-group comparisons for RNA-seq studies. BMC Genomics. 2020;21(1):75.

17. Flask [Available from: https://github.com/pallets/flask.

18. Lab S. Satija Lab: Seurat [Available from: https://satijalab.org/seurat/.

19. Marini F. Accessing the Human Cell Atlas Datasets. [Available from: https://bioconductor.org/packages/devel/data/experiment/vignettes/HCAData/inst/doc/hcadata.html.

20. Cudre-Mauroux P KH, Lim K-T, Rogers J, Simakov R, Soroush E, et al., editor A demonstration of SciDB: a science-oriented DBMS. Proc VLDB Endow; 2009.

21. Regev A. Human Cell Atlas Data Portal: Census of Immune Cells [Available from: https://data.humancellatlas.org/explore/projects/cc95ff89-2e68-4a08-a234-480eca21ce79.

22. Hafemeister C, Satija R. Normalization and variance stabilization of single-cell RNA-seq data using regularized negative binomial regression. Genome Biol. 2019;20(1):296.

23. Isaac Virshup GE, Sergei Rybakov, Fidel Ramirez, Tom White, Philipp Angerer, Alex Wolf, and Fabian Theis. Scanpy – Single-Cell Analysis in Python. [Available from: https://scanpy.readthedocs.io/en/stable/.

24. Philipp Angerer AW, Isaac Virshup, Sergei Rybakov. Anndata – Annotated Data. https://anndata.readthedocs.io/en/stable/. Accessed 07 June 2020 [Available from: https://anndata.readthedocs.io/en/stable/.

25. community TS. SciPy [Available from: https://docs.scipy.org/doc/scipy/reference/generated/scipy.io.mmwrite.html.

26. NCBI. Retrieve records annotated with a given biological feature 2007 [Available from: https://www.ncbi.nlm.nih.gov/Class/NAWBIS/Modules/InfoHubs/Exercises/infohubs_qa_biological_feature.html.

27. Morgan JI, Curran T. Stimulus-transcription coupling in the nervous system: involvement of the inducible proto-oncogenes fos and jun. Annu Rev Neurosci. 1991;14:421–51.

28. Stark R, Grzelak M, Hadfield J. RNA sequencing: the teenage years. Nat Rev Genet. 2019;20(11):631–56.

29. Srivatsan SR, McFaline-Figueroa JL, Ramani V, Saunders L, Cao J, Packer J, et al. Massively multiplex chemical transcriptomics at single-cell resolution. Science. 2020;367(6473):45–51.

30. Stoeckius M, Hafemeister C, Stephenson W, Houck-Loomis B, Chattopadhyay PK, Swerdlow H, et al. Simultaneous epitope and transcriptome measurement in single cells. Nat Methods. 2017;14(9):865–8.

31. Lacar B, Linker SB, Jaeger BN, Krishnaswami SR, Barron JJ, Kelder MJE, et al. Nuclear RNA-seq of single neurons reveals molecular signatures of activation. Nat Commun. 2016;7:11022.

32. Hoffmann M, Kleine-Weber H, Schroeder S, Kruger N, Herrler T, Erichsen S, et al. SARS-CoV-2 Cell Entry Depends on ACE2 and TMPRSS2 and Is Blocked by a Clinically Proven Protease Inhibitor. Cell. 2020;181(2):271–80 e8.

33. Zang R, Gomez Castro MF, McCune BT, Zeng Q, Rothlauf PW, Sonnek NM, et al. TMPRSS2 and TMPRSS4 promote SARS-CoV-2 infection of human small intestinal enterocytes. Sci Immunol. 2020;5(47).

34. Zhao G, Jiang Y, Qiu H, Gao T, Zeng Y, Guo Y, et al. Multi-Organ Damage in Human Dipeptidyl Peptidase 4 Transgenic Mice Infected with Middle East Respiratory Syndrome-Coronavirus. PLoS One. 2015;10(12):e0145561.

35. Zou X, Chen K, Zou J, Han P, Hao J, Han Z. Single-cell RNA-seq data analysis on the receptor ACE2 expression reveals the potential risk of different human organs vulnerable to 2019-nCoV infection. Front Med. 2020;14(2):185–92.

36. Qi F, Qian S, Zhang S, Zhang Z. Single cell RNA sequencing of 13 human tissues identify cell types and receptors of human coronaviruses. Biochem Biophys Res Commun. 2020;526(1):135–40.

37. Zhao Y ZZ, Wang Y, Zhou Y, Ma Y, Zuo W.. Single-cell RNA expression profiling of ACE2, the receptor of SARS-CoV. 2020.

